# Transmission and Detection of 0.1-2.5 THz Through Porcine Tympanic Membrane

**DOI:** 10.1101/2023.08.10.552875

**Authors:** Reza Shams, Zoltan Vilagosh, David Sly

## Abstract

Research has shown that exposure to high power THz radiation can cause thermal damage to the ear, leading to hearing loss and damage to the tympanic membrane. However, more research is needed to fully understand the effects of low intensity THz radiation on the ear and to determine safe exposure levels. This study investigates the transmission of 0.1 to 2.5 THz electromagnetic waves through porcine tympanic membrane samples. Similar to human tympanic membrane, porcine ear drum is a thin layer of tissue that separates the external ear from the middle ear and plays a crucial role in the process of hearing. Using THz time-domain spectroscopy, transmission of THz waves through ex vivo porcine tympanic membrane samples was measured. Results indicate that transmission of THz waves through the tympanic membrane is frequency dependent, with higher transmission observed at lower frequencies (0.1 to 0.5 THz) and lower transmission observed at higher frequencies (2 to 2.5 THz). This study provides new insights into the transmission of THz waves through the tympanic membrane and has potential to examine potential bioeffects as a result of THz interaction.

## 8.2. Introduction

While THz technology has been widely studied for its potential in a wide range of applications such as medical imaging and sensing (Fitzgerald et al., 2002; Taylor et al., 2011; Yan et al., 2022), its bioeffects on hearing and the tympanic membrane are not well understood. The tympanic membrane, also known as the eardrum, is a thin layer of tissue that separates the external ear from the middle ear and plays a crucial role in the process of hearing. It is responsible for transmitting sound vibrations from the air to the inner ear, where they are converted into electrical signals that are sent to the brain (Moller, 2012, pp.3-24).

The potential thermal effects of Terahertz (THz) radiation on the tympanic membrane are linked to the tissue’s absorption of energy, which consequently elevates its temperature. The absorption of energy from THz radiation by a tissue can trigger an increase in temperature, potentially compromising the structural integrity of the biological material. Both modeling and experimental studies have demonstrated that the threshold for most biological tissues is around 7.16 W/cm² (Dalzell et al., 2010). Concerning the tympanic membrane, non-radiation sources such as extended periods of high temperature and changes in temperature have been shown to alter its mechanical properties, ultimately leading to hearing loss (Hellmund et al., 1990). In general, a temperature rise of more than a few degrees Celsius can cause damage to the structures of the hearing system. Research has indicated that temperature changes in the middle ear are associated with a reduction in auditory nerve stimulation, damage to cochlear outer hair cells, and a decline in cochlear response (Gulick & Cutt, 1960; Noyes et al., 1996). Additionally, the non-thermal effects of THz radiation on biological tissues and specifically on the tympanic membrane, though not conclusively proven, are theorized to be connected to the interaction of the electromagnetic field with the biomolecules composing the tissue. These interactions are thought to be mediated by the thermal effect.

The degree of Terahertz (THz) absorption is contingent upon various factors, including the frequency, intensity, and duration of exposure, as well as the inherent properties of the tissue (Pickwell et al., 2004; Vilagosh et al., 2019; Nikitkina et al., 2021). Additionally, blood flow to the ear may influence the thermal response of the tympanic membrane (Meiners & Dabbs, 1977; Brinnel & Cabanac, 1989). Exposure to high-intensity THz radiation over a prolonged period can cause significant thermal damage to biological tissues, such as the tympanic membrane and adjacent tissues. This can lead to structural changes, potentially resulting in hearing loss. Biological tissues demonstrate a peak absorption coefficient at approximately 0.1 THz frequency. This is due to the interaction with water molecules within the tissues, while the duration and intensity of exposure can potentially amplify this effect (Cherkasova & Serdyukov, 2021; Hough et al., 2021).

The absorption of Terahertz (THz) radiation by tissues, including the skin and tympanic membrane, is a complex process that can be partially explained using the Beer-Lambert law. This law establishes a relationship between the intensity of radiation passing through a material, the amount of material it passes through, and the material’s absorption coefficient. For tissues and THz radiation, the absorption coefficient is a function of the radiation’s frequency and the tissue’s physical characteristics, such as its thickness and water content.

**Equation 1**

*Beer-Lambert’s Absorption Coefficient Equation*.

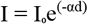

where I is the intensity of the THz radiation after passing through a distance d of skin, I0 is the initial intensity of the radiation, and α is the absorption coefficient of the skin at the frequency of the THz radiation. The absorption coefficient is typically expressed in units of cm^−1^.

However, it’s essential to consider that this equation is a simplification of the underlying complexities. Factors like the radiation’s frequency and intensity, the skin’s temperature, and its hydration state influence the absorption of THz radiation. Therefore, real-world predictions may vary.

The thermal effect ensuing from THz absorption is strongly associated with the electromagnetic (EM) radiation’s power density. Power density represents the power delivered by the EM radiation to a material per unit area. Higher power densities correspond to a more substantial deposition of energy into the material, resulting in more considerable thermal effects (Li et al., 2019). To measure the thermal effect of THz exposure, a simplified version of Pennes’ (1948) bioheat equation can be used. This equation provides an approximation, as it leaves out several influential factors like blood perfusion rate and blood’s heat capacity. However, since there are only limited number of capillaries only in the outer epithelial layer of the tympanic membrane (Hellstrom et al., 2003), their influence remains negligible and therefore can be approximated the simplified version.

**Equation 2**

*Simplified Bioheat Equation*

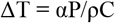

where ΔT is the temperature rise, α is the absorption coefficient of the material, P is the power density of the EM radiation, ρ is the density of the material, and C is its specific heat capacity.

This equation shows that the temperature rises in a material due to EM absorption is proportional to the power density and the absorption coefficient of the material. The thermal effect is also influenced by the density and specific heat capacity of the material, which determine how much energy is required to raise its temperature.

However, the relationship between power density and thermal effect isn’t linear. Higher power densities can trigger non-linear thermal effects, such as thermal runaway or thermal damage. Therefore, controlling the power density of EM radiation is crucial to prevent harmful thermal effects. Additionally, it’s important to note that the bioheat equation presumes homogeneity of the material, which isn’t the case with the tympanic membrane. As a result, this equation can provide an approximation at best.

A potential consequence of THz radiation’s thermal effect pertains to its impact on the ear’s transient receptor potential (TRP) channels, specifically within the tympanic membrane. TRP channels are protein structures that act as sensitive sensors, responding to shifts in temperature and pressure. When activated, they allow the ear to adapt to these changes (Romanenko et al., 2017). However, THz radiation can stimulate these TRP channels, potentially causing them to function abnormally. Such disruptions could lead to atypical responses, potentially compromising the tympanic membrane’s integrity and increasing the risk of hearing complications (Romanenko et al., 2017).

In addition to thermal effects, THz radiation may have non-thermal effects on the tympanic membrane and hearing. These effects may be the result of changes in the molecular structure of the tissue or changes in the physiology of the cells. One potential non-thermal effect of THz radiation on the tympanic membrane is the creation of reactive oxygen species (ROS), which can cause damage to the membrane’s structure and lead to hearing loss. ROS are highly reactive molecules that contain oxygen and are produced by the cells as a byproduct of normal cellular metabolism but can also be generated by external factors such as radiation, toxins, and pollutants (Finkel & Holbrook, 2000). ROS can cause damage to various biomolecules including DNA, proteins and lipids, if not properly neutralized by the body’s antioxidant defense system (Sitnikov et al., 2021). Studies have also suggested that THz radiation may cause changes in the expression of genes related to hearing and the function of the inner ear, which could potentially lead to hearing loss..

It’s important to note that the non-thermal effects of THz radiation on the ear are not well understood, and more research is needed to fully understand the mechanisms behind these effects, including the potential role of reactive oxygen species (ROS) and to determine safe exposure levels for human. In contrast, the thermal effects of THz radiation on the ear are better understood, and it is well established that this type of exposure has the potential to affect biological tissues, including the tympanic membrane, causing damage and potentially leading to hearing loss. Therefore, the aim of this study is to examine the propagation, transmission and intensity of THz radiation in order to better understand the thermal effects of THz radiation on the ear and to determine safe exposure levels for human.

## Method

The sample for this THz exposure study consists of two female porcine ears (total 4 ear samples), which were acquired from a butcher on the day of the study to ensure that the samples were fresh. The use of porcine ears in this study is a common practice in THz radiation research as the size and structure of the porcine ear is similar to that of the human ear. Using fresh samples ensures that the ears are in optimal condition for the study and allows for accurate measurements of the effects of THz radiation on the ear. The two female porcine ears were chosen to add a control aspect to the study. Porcine ears are considered a good model for human ear due to their morphological similarities, and their use allows for a better understanding of the effects of THz radiation on the human ear.

Ethics clearance had been obtained from the university for the use of scavenged tissue. This was done after a thorough review of the study’s design and protocols by the university’s ethics committee. The study was conducted in compliance with the regulations and guidelines set forth by the university ethics committee and safe transportation of biological materials. The tissue used in this study was obtained from an abattoir and was fresh on the day of the use, minimizing the time between the death of the pig and the start of the experiment. All efforts were made to ensure that no animal was killed or harmed for the purpose of this study and only used scavenged sample to ensure that the study was conducted in a humane manner. All necessary steps had been taken to ensure that the study met the ethical standards for research involving animals.

The BATOP THZ-TDS (Terahertz Time-Domain Spectroscopy) equipment is used in this study to measure the intensity, propagation, and transmission of THz radiation in the frequency range of 0.1 to 2.5 THz. THz-TDS is a non-destructive, non-contact method of measuring THz radiation that utilizes a femtosecond laser to generate THz pulses and a detector to measure the amplitude and phase of the THz radiation. This equipment allows for high-resolution measurements of THz radiation in the time domain and can be used to measure the absorption and transmission of THz radiation in various materials. The equipment is a highly advanced and reliable system and is capable of measuring THz radiation in a wide range of frequencies and intensities, and the ability to measure the frequency range of 0.1 to 5 THz but is more accurate at 0.1 to 2.5 THz and is particularly useful for this study.

The dissection process involved the careful removal of the external ear and pinna, as well as the ear canal and the middle ear structures, including the tympanic membrane, in order to expose them to the radiation. This dissection process was carried out under sterile conditions to prevent contamination of the sample. The dissection was performed as quickly as possible to minimize the time between the death of the pig which was acquired on the day from an abattoir and the start of the experiment, in order to ensure that the sample was as fresh as possible. Once the dissection was complete, the tympanic membrane was carefully handled to avoid damage, and the sample was immediately placed in the 3D printed sample holder and inserted into the sample holder of the BATOP THZ-TDS equipment for exposure to THz radiation.

## Results

The study found that the transmission of THz radiation through porcine ears is frequency-dependent, with higher transmission at lower frequencies and lower transmission at higher frequencies. The absorption of THz radiation is higher at higher frequencies, leading to lower penetration of the radiation. Tissues with higher water content had reduced transmission of THz radiation and therefore increased absorption, with muscle having the highest absorption coefficient, followed by brain, tympanic membrane, and skin. These results are consistent with previous studies that have shown that biological tissues have a higher absorption coefficient at higher frequencies, and a higher transmission at lower frequencies. The findings of this study suggest that the transmission and absorption of THz radiation through biological tissues, including the tympanic membrane, is frequency-dependent and is affected by the water content of the tissue.

The results of the study indicate that the transmission of THz radiation through biological tissues, including the tympanic membrane, is influenced by the frequency of the radiation and the water content of the tissue. The study’s finding that the transmission of THz radiation is higher at lower frequencies and lower at higher frequencies is consistent with previous research that has shown that the attenuation of THz radiation in biological tissues increases with frequency. The study’s finding that tissues with higher water content, including many of the tissues found in the ear, are more likely to absorb THz radiation, leading to a higher risk of thermal damage, is also consistent with previous research. The study’s results provide important insight into the potential effects of THz radiation on biological tissues, including the ear.

Assuming compliance with the prevailing safety guideline that limits radiation exposure to 0.2mW/cm², the resultant increase in temperature due to the absorption of 0.2 milliwatts of radiation of tissue is influenced by a multitude of variables. This includes the specific heat capacity of the tympanic membrane, which stands at approximately 3500 J(kg.K), and its density, measured at 1.2 x 103 kg/m^3^. Furthermore, the duration of radiation exposure and the fraction of the radiation that is successfully absorbed also play significant roles. The complex interaction between these parameters fundamentally dictated the thermal behavior of biological tissues subjected to THz radiation exposure.

The thermal effects of terahertz (THz) radiation exposure were computed for muscle, tympanic membrane (TM), and skin across a range of frequencies from 0.1 THz to 2.0 THz, employing the simplified bioheat equation. The absorption coefficients, tissue densities, and specific heat capacities were used in the calculations. According to the findings, the temperature change, which directly corresponds to thermal effects, varies across tissue types and THz frequencies. Muscle demonstrated a temperature change ranging from 0.034°C to 0.098°C, TM exhibited a range of 0.028°C to 0.132°C, while skin presented a change in the range of 0.063°C to 0.200°C. Among all tissues, the skin demonstrated the highest sensitivity to THz radiation across the analyzed frequencies.

As shown in Figure 3, comparing current occupational limit power density of 10 W/m² (equivalent to 1 mW/cm²) as opposed to the general population limit of 2 W/m². The results demonstrated a marked increase in the temperature change across all tissues and frequencies compared to the lower power density. For instance, at a frequency of 1 THz, the temperature rise in muscle tissue was approximately 0.452°C at 10 W/m², an increase of around 4.5 times compared to the 0.1°C rise observed at 2 W/m². Similarly, the temperature change in the tympanic membrane at the same frequency rose from 0.067°C to 0.255°C, and the skin from 0.077°C to 0.350°C. This underlines the direct proportionality between power density and thermal effect, signifying that a five-fold increase in power density translates into a roughly five-fold elevation in thermal effect, assuming other factors like absorption coefficient, tissue density, and specific heat capacity remain constant.

**Figure 1.**
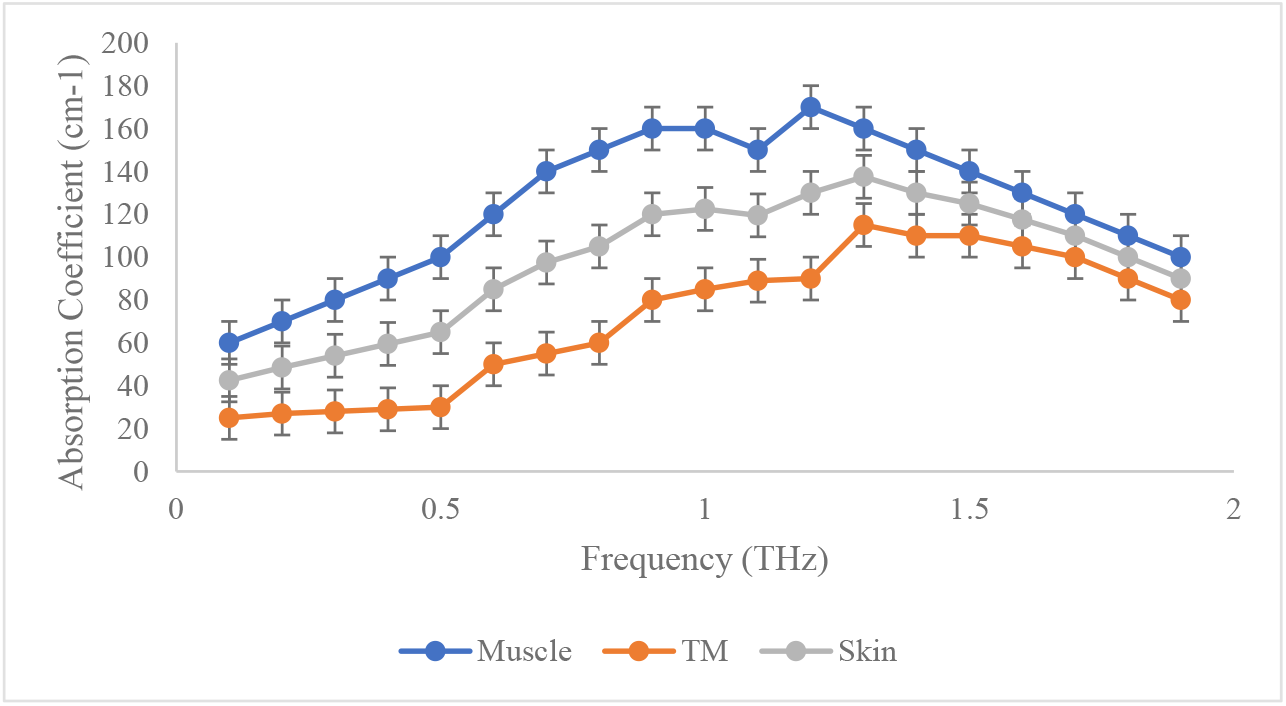
Absorption Coefficient of Pig Skin, Muscle and Tympanic Membrane (TM) at 0.1-2 THz. *Note*. Muscle (porcine tissue), TM (tympanic membrane and Skin (porcine tissue).

**Figure 2.**
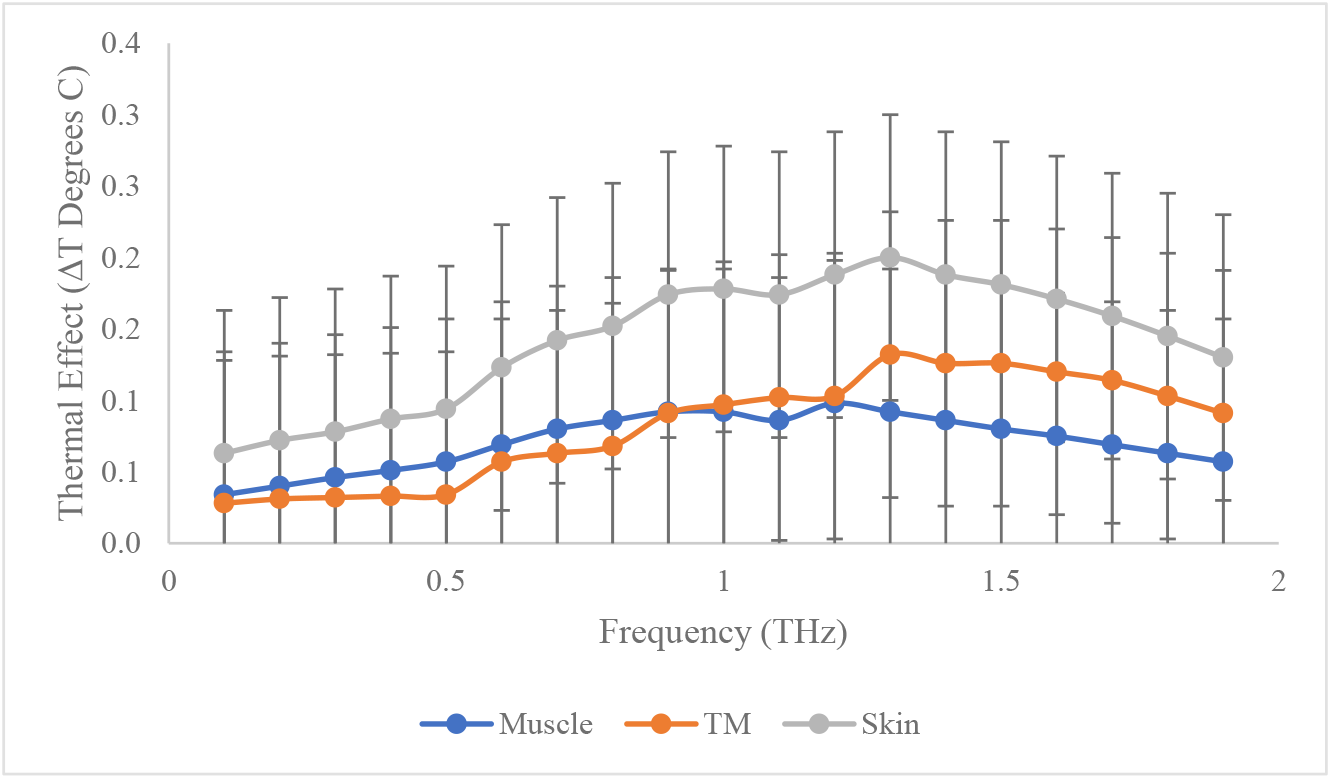
Calculated Thermal Effects of 2 w/m^2^ THz Exposure in Pig Skin, Muscle and Tympanic Membrane (TM) at 0.1-2 THz. *Note*. Muscle (porcine tissue), TM (tympanic membrane and Skin (porcine tissue).

**Figure 3.**
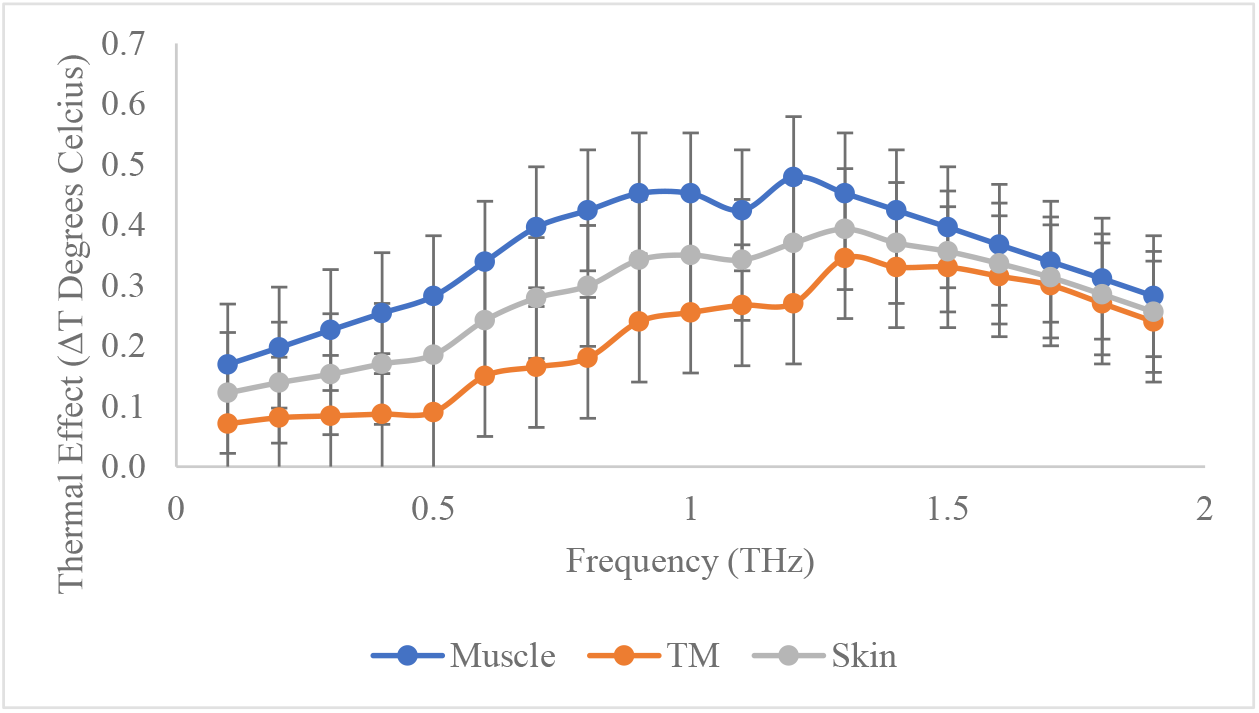
Calculated Thermal Effects of 10 w/m^2^ THz Exposure in Pig Skin, Muscle and Tympanic Membrane (TM) at 0.1-2 THz. *Note*. Muscle (porcine tissue), TM (tympanic membrane and Skin (porcine tissue).

## Discussion

The outcomes of this investigation offer valuable perspectives on how THz radiation traverses and is absorbed by various biological tissues, including the sensitive tympanic membrane. The observation that the transmission of THz radiation through these tissues hinges on frequency aligns with preceding studies that have reported escalating attenuation of this radiation type as frequency increases. This pivotal finding underscores the notion that transmission is enhanced at lower frequencies while being diminished at higher ones, bearing significant implications for establishing safe exposure parameters. Given this, these findings propose that the prevailing safety thresholds may engender differing thermal effects, contingent on the intensity applied.

The results of this study are pivotal for evaluating secure exposure guidelines for THz radiation, particularly considering future telecommunication, as well as medical applications like imaging and sensing. These data implies that the absorption of THz radiation amplifies with increased frequencies, leading to a reduction in the radiation’s penetration. This signals that a high intensity and sustained exposure to THz radiation may provoke significant thermal damage to the tympanic membrane and surrounding tissues, triggering structural changes and potentially inducing hearing loss. This assertion is particularly salient in light of the occupational limit and the resultant temperature rise of 0.452°C at 10 W/m². Factors such as extended exposure could potentially intensify these effects further.

Moreover, the results demonstrating that tissues with a greater water concentration manifest decreased transmission yet heightened absorption of THz radiation correspond with existing research. The amplified absorption coefficient of THz radiation in tissues rich in water can be attributed to the interplay between the radiation and the water molecules embedded in the tissue. Thus, tissues with a higher water content may be more prone to thermal damage from THz radiation exposure. Ultimately, this study suggests that the probable thermal impacts of THz radiation on biological tissues, including the tympanic membrane and auditory mechanism, are governed by distinct tissue properties, primarily its water content.

However, it should be emphasized that the real-world effects of temperature on biological tissues extend beyond our theoretical computation. Biological tissues frequently experience temperature losses due to an array of elements such as ambient temperature, blood circulation, and other physiological phenomena. Environmental temperature can exert a substantial impact on bodily and surrounding tissue temperatures. In colder surroundings, heat is dissipated into the environment, leading to a drop in tissue temperature, while warmer surroundings could elevate tissue temperatures through heat absorption. Blood supply also significantly influences tissue temperature; efficient blood flow aids in heat distribution within the body, whereas reduced blood flow to specific regions can induce localized cooling. All these factors synergistically influence changes in tissue temperature, with wide-ranging implications on THz interaction and subsequent thermal effect.

THz radiation exposure poses particular concerns for individuals susceptible to thermal effects, especially in hotter climates where ambient temperatures are elevated. The thermal effects of THz radiation can provoke localized temperature spikes in soft tissues, potentially causing damage if the temperature rise is substantial. In extreme scenarios, exposure to high levels of THz radiation may induce burns or other thermal injuries, particularly if the body’s natural mechanisms fail to adequately disperse or absorb the radiation. Therefore, careful consideration of the potential risks associated with THz radiation and the implementation of precautions to minimize exposure are essential when engaging with this radiation type.

Understanding the non-thermal effects of THz radiation on biological tissues, including the ear, remains a challenge and necessitates further investigation to completely decipher these mechanisms. Yet, it’s apparent from these findings that the thermal consequences of THz radiation on biological tissues, specifically the tympanic membrane and the auditory system, are more readily understood. It’s widely accepted that this form of exposure can affect biological tissues, leading to damage and potentially precipitating hearing loss. Therefore, examining the potential moderating influence of thermal effect on non-thermal impact demands additional research for a comprehensive understanding of the potential risks associated with exposure to THz radiation.

## Reference

Brinnel, H., & Cabanac, M. (1989). Tympanic temperature is a core temperature in humans. Journal of Thermal Biology, 14(1), 47–53.

Cherkasova, O. P., Serdyukov, D. S., Nemova, E. F., Ratushnyak, A. S., Kucheryavenko, A. S., Dolganova, I. N., … & Tuchin, V. V. (2021). Cellular effects of terahertz waves. Journal of Biomedical Optics, 26(9), 090902–090902. https://doi.org/10.1117/1.JBO.26.9.090902

Dalzell, D. R., McQuade, J., Vincelette, R., Ibey, B., Payne, J., Thomas, R., … & Wilmink, G. J. (2010, February). Damage thresholds for terahertz radiation. In Optical Interactions with Tissues and Cells XXI(Vol. 7562, pp. 149–156). SPIE. https://doi.org/10.1117/12.849243

Finkel, T., & Holbrook, N. J. (2000). Oxidants, oxidative stress and the biology of ageing. Nature, 408(6809), 239–247.

Fitzgerald, A. J., Berry, E., Zinov’ev, N. N., Homer-Vanniasinkam, S., Miles, R. E., Chamberlain, J. M., & Smith, M. A. (2003). Catalogue of human tissue optical properties at terahertz frequencies. J Biol Phys, 29(2-3), 123–128. https://doi.org/10.1023/a:1024428406218

Gulick, W. L., & Cutt, R. A. (1960). III The Effects of Abnormal Body Temperature upon the Ear: Cooling. Annals of Otology, Rhinology & Laryngology, 69(1), 35–50.

Hellmund, S., Begall, K., & Preibisch-Effenberger, R. (1990). Meningitis and hearing damage in children. Padiatrie und Grenzgebiete, 29(1), 13–17.

Hough, C. M., Purschke, D. N., Huang, C., Titova, L. V., Kovalchuk, O. V., Warkentin, B. J., & Hegmann, F. A. (2021). Intense terahertz pulses inhibit Ras signaling and other cancer-associated signaling pathways in human skin tissue models. Journal of Physics: Photonics, 3(3), 034004. https://doi.org/10.1088/2515-7647/abf742

Li, K., Sasaki, K., Watanabe, S., & Shirai, H. (2019). Relationship between power density and surface temperature elevation for human skin exposure to electromagnetic waves with oblique incidence angle from 6 GHz to 1 THz. Physics in Medicine & Biology, 64(6), 065016. https://doi.org/10.1088/1361-6560/ab057a

Meiners, M. L., & Dabbs Jr, J. M. (1977). Ear temperature and brain blood flow: Laterality effects. Bulletin of the Psychonomic society, 10(3), 194–196.

Møller, A. R. (2012). Hearing: anatomy, physiology, and disorders of the auditory system. Plural Publishing.

Nikitkina, A. I., Bikmulina, P. Y., Gafarova, E. R., Kosheleva, N. V., Efremov, Y. M., Bezrukov, E. A., … & Timashev, P. S. (2021). Terahertz radiation and the skin: a review. Journal of Biomedical Optics, 26(4), 043005–043005. https://doi.org/10.1117/1.JBO.26.4.043005

Noyes, W. S., McCaffrey, T. V., Fabry, D. A., Robinette, M. S., & Suman, V. J. (1996). Effect of temperature elevation on rabbit cochlear function as measured by distortion-product otoacoustic emissions. Otolaryngology—Head and Neck Surgery, 115(6), 548–552.

Pennes, H. H. (1948). The thermal conductivity of living tissue. Journal of Applied Physiology, 1(2), 93–122.

Romanenko, S., Begley, R., Harvey, A. R., Hool, L., & Wallace, V. P. (2017). The interaction between electromagnetic fields at megahertz, gigahertz and terahertz frequencies with cells, tissues and organisms: risks and potential. Journal of The Royal Society Interface, 14(137), 20170585. https://doi.org/10.1098/rsif.2017.0585

Sitnikov, D. S., Ilina, I. V., Revkova, V. A., Rodionov, S. A., Gurova, S. A., Shatalova, R. O., … & Baklaushev, V. P. (2021). Effects of high intensity non-ionizing terahertz radiation on human skin fibroblasts. Biomedical optics express, 12(11), 7122–7138. https://doi.org/10.1364/BOE.440460

Taylor, Z. D., Singh, R. S., Bennett, D. B., Tewari, P., Kealey, C. P., Bajwa, N., … & Grundfest, W. S. (2011). THz medical imaging: in vivo hydration sensing. IEEE transactions on terahertz science and technology, 1(1), 201–219. https://doi.org/10.1109/TTHZ.2011.2159551

Vilagosh, Z., Lajevardipour, A., & Wood, A. (2019). An empirical formula for temperature adjustment of complex permittivity of human skin in the terahertz frequencies. Bioelectromagnetics, 40(1), 74–79. https://doi.org/10.1002/bem.22156

Yan, Z., Zhu, L. G., Meng, K., Huang, W., & Shi, Q. (2022). THz medical imaging: from in vitro to in vivo. Trends in Biotechnology, 40(7), 816–830. https://doi.org/10.1016/j.tibtech.2021.12.002

